# Increased host diversity limits bacterial generalism but may promote microbe-microbe interactions

**DOI:** 10.1101/2024.04.24.590977

**Authors:** Iris A Holmes, José G Martínez-Fonseca, Rudolf von May, Briana A Sealey, Peter A Cerda, Maggie R Grundler, Erin P Westeen, Daniel Nondorf, Joanna G Larson, Christopher R Myers, Tory A Hendry

**Affiliations:** Cornell Institute of Host-Microbe Interactions and Disease, Cornell University, Ithaca, NY 14853, USA; Department of Microbiology, Cornell University, Ithaca, NY 14853, USA; Department of Public and Ecosystem Health, Ithaca, NY 14853, USA; Museum of Zoology and Department of Ecology and Evolutionary Biology, University of Michigan, Ann Arbor, MI 48109, USA; School of Forestry, Northern Arizona University, Flagstaff, AZ 86011, USA; Biology Program, California State University Channel Islands, Camarillo, CA 93012, USA; Department of Integrative Biology, University of Texas, Austin, TX 78712, USA; Department of Environmental Science, Policy, and Management and Museum of Vertebrate Zoology, University of California, Berkeley, CA 94720, USA; Department of Biology, University of Virginia, Charlottesville, VA 22903, USA; Department of Biological Sciences and Museum of Biodiversity, University of Notre Dame, Notre Dame, IN 46556, USA; Center for Computing & Laboratory of Atomic and Solid State Physics, Cornell University, Ithaca, NY 14853, USA

## Abstract

Host-associated bacteria vary in their host breadth, which can impact ecological interactions. By colonizing diverse hosts, host generalists can have disproportionate ecological impacts. For bacteria, host generalism may advantageous, particularly when the availability of specific hosts is variable. It is unclear how much the ability to evolve generalism, by inhabiting diverse hosts, is constrained in host-associated bacteria. We hypothesized that constraints on bacterial generalism will differ depending on the availability of specific host species. To test this, we assessed patterns of diversity and specialization in the cloacal microbiomes of reptile communities from the temperate zone to the tropics, where the diversity and abundance of host species varies substantially. Within these communities, generalist taxa tended to be Proteobacteria, whereas specialists tended to be Firmicutes. We found that bacterial generalists were less prevalent in the highest diversity host communities, and in keeping with this, Proteobacteria were less diverse in these communities. Generalist taxa became relatively less widespread across host species only in the two most diverse host communities. We therefore conclude that the constraint on generalism is not driven by absolute incompatibility with some host species, but rather from competition with host adapted specialist lineages. In the high-diversity communities, we found that the successful generalists, typically Proteobacteria, were disproportionately likely to co-occur with one another across evolutionarily disparate hosts within the community. Our data indicate that bacterial lineages can adapt to the evolutionary pressures of high diversity host communities either by specializing on hosts or by forming cohorts of co-occurring bacterial lineages.

## Introduction

Like all organisms, bacterial lineages experience conflicting selective pressures between specialization and generalism (Bell and Bell 2020). When resources are consistently available, specialists have a competitive advantage relative to generalists, whereas in fluctuating environments generalists have a survival advantage (Bono et al. 2017; Chen et al. 2021). While most bacterial lineages are habitat specialists, (Mariadassou et al. 2015; Sriswasdi et al. 2017), host-associated generalist lineages can carry outsized ecological importance as pathogens (Aijuka and Buys 2019) or by contributing beneficial metabolic processes to hosts (Sriswasdi et al. 2017; Scott et al. 2020). Testing patterns of bacterial generalism across host communities with different degrees of host phylogenetic diversity and species richness can help to identify absolute and relative ecological constraints on bacteria niche breadth, but such studies are relatively rare (Cobian et al. 2019). To address this gap, we analyzed patterns of host generalism and ecological interactions in mucosal microbiome communities across vertebrate hosts in communities that span an order of magnitude change in diversity. Vertebrate mucosal microbiomes tend to have tens to hundreds of bacterial lineages, acquired from both horizontal transmission and vertical inheritance (Van Veelen et al. 2017; Ambrosini et al. 2019; Bodawatta et al. 2021; Hernandez et al. 2021). Individual lineages vary from highly host-specific (Eren et al. 2015; Mallott and Amato 2021) to surviving in virtually any host species as well as outside of hosts (Bell and Bell 2020), making vertebrate mucosal microbiomes ideal communities in which to identify the selective pressures shaping bacterial host generalism.

We sampled the functionally important cloacal microbiome of snakes and lizards. Reptile cloacas are semi-oxygenated habitats at the terminus of the digestive, urinary, and reproductive tracts (Grond et al. 2018). Cloacal microbiota composition can be impacted by host diet, habitat and stress levels (Colston 2017; Ambrosini et al. 2019; Van Veelen et al. 2020). These communities can include parasites that lead to morbidity or mortality (Johne et al. 2002; Styles et al. 2004; Curtiss et al. 2015; Tillis et al. 2021), exerting selective pressure on hosts. The cloaca also houses microbial lineages that inoculate the eggshell during laying and provide protection against fungal infection, another source of selection on host control of the microbiome (Bunker et al. 2021). This egg inoculation could also provide opportunity for vertical transmission of microbes (Li et al. 2022; Bunker and Weiss 2024).

Gut bacteria can colonize the cloacal microbiome, but bacterial lineages also colonize through mating or through contact with the external environment (Colston 2017; Kohl et al. 2017; Hoffbeck et al. 2023). Adaptation to the cloacal habitat, competition with the existing community and interaction with host immune processes may all structure the cloacal community (Bolnick et al. 2014; Hansen et al. 2015; Holmes et al. 2017; Verster and Borenstein 2018). The potential combination of pathogenic, commensal, and mutualistic lineages, as well as both horizontal and vertical transmission, makes the cloacal microbiome an ideal site for studying host-associated microbial community ecology.

Proteobacteria and Firmicutes are common phyla in many vertebrate gut microbiomes (Ley et al. 2008; Youngblut et al. 2019; Song et al. 2020), including in the reptile cloaca (Colston and Jackson 2016; Hoffbeck et al. 2023). Broadly, these two groups typify two different evolutionary strategies. Proteobacteria include extreme host generalists and are generally horizontally transmitted across hosts (Gripp et al. 2011; Dandekar et al. 2012; Touchon et al. 2020). Some Firmicutes lineages can be more host-specific, can be inherited vertically (Moeller et al. 2018), and have been shown to codiversify with their hosts (Moeller et al. 2016; Sanders et al. 2023). We therefore hypothesize that these two taxa may respond differently to selective pressures for specialization or generalism.

Vertebrate species diversity increases tenfold from the temperate zone to the tropics (Lawrence and Fraser 2020). Microbes that specialize on a narrow range of host phylogenetic diversity may therefore have more difficulty finding appropriate new host individuals via horizontal transmission within a more diverse host community. Vertical transmission may be more effective in this context. Conversely, generalist microbes would need to be able to adapt to a greater number of selective pressures across diverse hosts, including different immune responses, nutrient availability, and competition from host specialist microbes (Woodcock et al. 2017; Mouftah et al. 2021; Maritan et al. 2024).

Here we test three fundamental hypotheses about the ecology and evolution of host generalism in host-associated bacterial lineages. First, we hypothesize that high-diversity host communities will have the highest diversity of microbial lineages. As the number of host lineages increase, we expect the number of specialist bacterial lineages to increase as well, as has been observed in host-specific parasite lineages (Bordes et al. 2011; Clark 2018). In further support of this idea, free-living generalist bacteria are more diverse toward the tropics (Zuo et al. 2023). Second, we hypothesize that microbes in high diversity host communities will either be specialized on specific host lineages or highly generalist (Sant et al. 2021), with fewer lineages showing midrange generalism. Finally, we hypothesize that host generalism will necessitate flexibility with respect to bacteria-bacteria interactions, and that we will therefore find stronger evidence of bacteria-bacteria interactions between host specialist lineages. Host specialization might promote mutualist or competitive interactions between bacterial lineages by repeatedly bringing together specialized microbial lineages within a host species. Using the outcomes of these three lines of inquiry, we will identify whether bacterial generalism in our sampled communities is constrained by bacterial lineages being incompatible with hosts or by more diffuse ecological interactions between the total host-microbiome ecosystem.

## Methods

### Sample collection

We collected samples from temperate forest habitats near Ann Arbor, MI and Girard, GA in the United States. We also collected in the Nicaraguan highlands at Las Brisas del Mogoton (montane pine forest and cloud forest), dry forest at Asosoca Lake, and tropical wet forest at Refugio Bartola. Further sampling detail is available in Martínez-Fonseca et al. 2024. In Peru, we sampled in tropical wet forest at Los Amigos Biological Station (Figure S1; Table 1; Table S1). We divided sites into host richness categories by the number of host families present, as host family consistently predicts pairwise microbiome distances whereas host species and genus do not (Hoffbeck et al. 2023). Sites fell out into low, medium, and high host family diversities (Table 1), with the temperate forest sites being lowest and the tropical wet forest sites highest. To quantify the diversity of host species and higher taxa, we used the IUCN ‘scaled reptile’ range polygons (IUCN 2023). We plotted them using the R package ‘sf’ (Pebesma 2018; Pebesma and Bivand 2023) and found the number of ranges that intersected the centroid points of our sampling locations (Table 1).

Because tropical seasonality differs from temperate seasons, we cannot directly compare sampling times. In the temperate zone, we sampled in spring, as our host species were actively foraging following winter (Table S1). In the tropics, we sampled at the start of the wet season (November in Peru, May in Nicaragua). Tropical reptiles do not all hibernate during the dry season, but they dramatically increase activity and feeding at the start of the wet season, leading to a similar set of behaviors as the temperate species display in spring.

We captured animals using hand grabs for lizards and snakes, or funnel trapping and lassoing with surgical silk for lizards (Lettink and Hare 2016). We searched during day and night using unstructured surveys. Techniques were approved by University of Michigan UCUCA Protocol #PRO00006234 and protocol # I16005 issued by Georgia Southern University IACUC to Christian L. Cox. We sampled microbiome DNA using a sterile rayon swab (MW113) inserted into the cloaca and quickly removed to collect cloacal mucosa and epithelial cells (Colston et al. 2015). Swabs were stored in RNAlater in the field and kept at ambient temperature away from direct sun for no more than five weeks. Samples were stored at -20C in the lab until DNA extraction.

### Laboratory methods

Total DNA was extracted from our swabs using Qiagen DNEasy blood and tissue kits. We incubated swabs for 12 hours, followed by vortexing prior to extraction. We sequenced the samples on a MiSeq platform with version 2 reagents at the University of Michigan Microbiome Core Facility, using primer sets developed by (Kozich et al. 2013) to amplify 252 bp from the 16S rRNA V4 region. Each sequencing run had a positive (mock community) and negative (water) control.

### Bioinformatics

To process our data, we used the qiime2 platform (Bolyen et al. 2019). We used the dada2 module to dereplicate, denoise, and filter out data for chimeras (Callahan et al. 2016). The algorithm clustered reads into amplicon sequence variants (ASVs). Amplicon sequence variants are the most specific evolutionary unit that can be inferred from amplicon data. We then built an insertion tree with the representative sequences for each ASV identified by the dada2 module (Callahan et al. 2016). We used the greengenes 13.8 database as a set of reference sequences (DeSantis et al. 2006). We processed our outputs using the qiime2R library embedded in a custom R script (Bisanz 2018). We used the node annotation from the insertion tree output to assign taxonomic positions to our ASVs. We also generated a host by ASV matrix, populated by coverage depth values. This post-processing was done using custom scripts in R v4.3.3 which will be available in a Zenodo repository on publication of this manuscript.

We filtered our results using a custom R script by removing any ASV with a coverage depth of five or fewer to account for index hopping (Van Der Valk et al. 2020). We then removed any host that had under 10,000 total reads across all ASVs. We chose this value as a compromise between coverage depth and the number of hosts we retained in our dataset (Figure S2). We removed any host with fewer than 50 called ASVs, as these results may represent sampling, extraction, or sequencing bias (Figure S3). Finally, we generated a reference dataset in which each location was subsampled to sixteen individuals to match our least deeply sampled location. For two locations, Georgia and the Nicaraguan highlands, we removed samples that were more than 5km distant from our main sampling location. We removed the samples to avoid artificially increasing location-wide metrics of diversity due to spatial variation. We removed all singleton ASVs, as we are primarily interested in interactions of ASVs with hosts and with each other. We then rarefied to 180 ASVs per location, to match the locality with the fewest ASVs. The six rarefied host x ASV matrices, which we refer to as our rarefied dataset, will also be available in our Zenodo repository.

### Phylum-level community structure and diversity

To visualize the relationship between within-location ASV host generalism and prevalence across hosts, we calculated host phylogenetic diversity for each ASV in our rarefied dataset using the ses.pd function from the R package ‘picante’ (Kembel et al. 2010). Negative values represented specialists and positive values represented generalists. The index compared the observed phylogeny of an ASV’s hosts to a size-matched random sample of hosts. We generated 9999 random host samples for each ASV. To calculate the index, we subtracted the mean phylogenetic distance between the null host subsamples from the distance between the real hosts, then divided by the standard deviation of the host distances from the null datasets. We could not calculate the metric for ASVs that were present in all hosts, because the standard deviations of their ‘null’ samples are zero. For these ASVs, we assigned the maximum value of the metric present in any other sample for that site. This approach underestimates the true maximum value of diversity, making it conservative with respect to the distinctions we test here.

To assess the predictors of microbiome similarity across hosts, we used a PERMANOVA test in the R package ‘vegan’ (Oksanen et al. 2018). We used location, host family, and a host family x location interaction term as explanatory variables. The PERMANOVA approach first found pairwise Bray-Curtis distances between hosts in ASV space. We used our non-rarefied dataset for this analysis. For each pre-determined group (i.e. hosts from a given location or host from a given family), the algorithm found the mean distances between individuals. For comparison, distances are found between equal numbers of randomly chosen individuals. If the mean distances between individuals within the predetermined groups were smaller than the mean distances between randomized groups, the explanatory variables may have biological significance. The proportion of randomized distances that are larger than the measured distance from the p-value. We used 99,999 permutations to find this value. We first tested all ASVs, then Proteobacteria ASVs only and Firmicutes ASVs only. To address the problem of multiple comparisons, we Bonferroni corrected our significance value for this paper to 0.0001. This is a conservative value, as it represents the total possible number of comparisons we could perform. For some of our later analyses, we use statistical tests that look for differences in values across all sites. If those tests are significant, we then assess pairwise comparisons between sites. To set our Bonferroni value, we found the total number of comparisons that we could perform if we were to do every pairwise comparison possible.

We hypothesized that high host diversity communities will have higher total ASV diversity, driven by host-specific lineages. To test this hypothesis, we looked at the total diversity of ASVs in our rarefied dataset. We plotted a community-wide species accumulation curve for each location using the ‘poolaccum’ command in the R package ‘vegan,’ first with all ASVs, then with only Proteobacteria and only Firmicutes ASVs. We used the first-order jackknife approach to account for the relatively small number of hosts in our subsamples (Smith and Van Belle 1984). To assess microbial diversity within hosts, rather than across the full community, we applied the Shannon diversity index implemented in the R package ‘vegan’ (Oksanen et al. 2018) and the phylogenetically informed Faith’s diversity index in the R package ‘picante’ to each host in our rarefied dataset (Kembel et al. 2010). Finally, we assessed specialization and generalism of ASVs in each location using the host diversity index from the ses.pd function (Procheş et al. 2006). For each diversity metric, we implemented an ANOVA test between locations using the function ‘aov’ in base R, first testing the full dataset, then only the Firmicutes and only the Proteobacteria lineages to capture differences in behavior between them. If the ANOVA was significant, we compared pairwise differences between sites using the ‘TukeyHSD’ function, also in base R.

### Comparing ASV interactions between host communities

If multiple microbial lineages all specialize on the same hosts, the lineages will have more opportunities for ecological interactions. We hypothesized that interactions between ASVs were stronger in host communities that include a greater proportion of specialist ASVs. To test this hypothesis, we found pairs of ASVs that occurred in the same hosts more or less frequently than would be expected by chance using the R package ‘cooccur’ (Griffith et al. 2016). Greater than expected overlap (positive co-occurrence) could indicate either shared host specializations or some type of direct interaction. Rarer than expected overlap (negative co-occurrence) could indicate either specialization on different hosts or competitive exclusion. The ‘cooccur’ package used an equation based on combinatorics to calculate the probability that pairs of ASVs both occupy the observed set of hosts, given each ASV’s prevalence in the dataset (Veech 2013). We identified pairs of ASVs with both negative and positive co-occurrence probabilities with a 0.05 threshold. Since the co-occurrence test can be sensitive to sampling effects, we subsampled our full dataset to 16 individuals and 140 ASVs per site, and repeated the subsampling 50 times. With each subsampled dataset, we generated a matching null dataset by randomizing the order of ASVs counts within columns to disrupt interactions between ASVs and hosts while retaining the underlying prevalence structure of the community. We ran co-occurrence analyses on the 50 matched subsampled and null datasets.

For each rarefaction run, we recorded the number of positively and negatively co-occurring ASV pairs per site at a significance value of 0.05. We also identified the prevalence of each interacting ASV across hosts. To test the hypothesis that shared host specificity drove co-occurrence, we calculated the Unifrac distance between the hosts of the ASVs in each pair using the R package ‘GUniFrac’ (Chen et al. 2023). Unifrac measures the amount of shared versus non-shared branch length on a phylogenetic tree between two samples (Lozupone and Knight 2005). We would expect smaller values for host specialists. We also calculated the patristic distance between ASVs in the co-occurring pairs. To test whether co-occurrence was driven by repeated within-host diversification, we calculated the mean nearest neighbor patristic distance and the overall mean patristic distance for all ASVs in each location in each run. If co-occurrence is driven by repeated within-host diversification, co-occurring pairs should have phylogenetic distances similar to this nearest-neighbor value. To account for nonrandom patterns of prevalence in our co-occurring ASVs, we generated a second null comparison by randomly sampling ASVs that matched the prevalence of each member of a pair and re-calculating the host Unifrac and ASV patristic distance metrics. For each descriptive metric, we performed an ANOVA to test for significant differences between sites, followed by pairwise site comparisons with a Tukey test if the ANOVA was significant. We tested for significant differences between real and null datasets within locations using t-tests. By inspection, we identified an excess of Proteobacteria-Proteobacteria pairs. To test whether this pattern was statistically significant, we found the proportion of all positively co-occurring pairs that were made up of two Proteobacteria ASVs in each run, and used a t-test to compare that value to the squared value of the proportion of total Proteobacteria ASVs in each corresponding subsampled dataset. This value represents the expected number of Proteobacteria-Proteobacteria pairs that would be drawn at random.

In addition to the pairwise co-occurrence values, we used a community detection algorithm to find modules of co-occurring ASVs within our localities. This approach allowed us to identify larger interacting units. We generated a further 50 matched rarefied and null datasets for this analysis. For each, we built a bipartite network in which each ASV was linked to the hosts in which it occurred. ASVs could only be associated with each other through their connections to the same hosts and vice versa. We calculated the modularity of the network, a statistic that describes the difference between the number of shared host connections between ASVs within modules compared to outside modules. Higher modularity values indicated stronger interactions. We used the ‘metaComputeModules’ command in the R package ‘bipartite’ (Dormann et al. 2008; Liu and Murata 2010; Dormann and Strauss 2014; Beckett 2016). We identified the number of modules in each run, and the number of hosts and ASVs in each module. We compared these values across sites using ANOVA, followed by pairwise Tukey tests for significant interactions, and compared real and randomized datasets within each location using t-tests.

We assessed the average phylogenetic clustering of hosts and ASVs in our real and randomized modules. We applied the ses.pd specialization index to the hosts of each module and found the mean value for the modules in each run. We found the mean of the mean pairwise distances between ASVs in the modules, standardized by the mean distance between ASVs in the rarefied dataset. Values less than one indicate phylogenetic clustering among ASVs in modules, while values greater than one indicate phylogenetic overdispersion.

## Results

### Bacterial diversity and generalism within and across communities

We found that our two most common bacterial phyla differed in their prevalence and specificity within and between sites. We identified a trend toward higher prevalence and host generalism in Proteobacteria compared to Firmicutes in our low and mid-diversity sites, with both phyla being more host specialized in our high-diversity locations (Figure 1). Our PERMANOVA showed that location, family, and location x family interaction terms were all significant for our full dataset (p < 1×10^-5^, R^2^ = 0.083, 0.101, and 0.086 respectively; Figure 2). Using Firmicutes ASVs only, host family is the only significant predictor variable (p = 1×10^-5^, R2 = 0.058). Conversely, the only significant predictor using Proteobacteria ASVs is location (p = 1×10^-5^, R2 = 0.0962). Accumulation curves of the number of unique ASVs recovered with the successive addition of increasing numbers of hosts showed similar levels of diversity across all locations when all ASVs in the rarefied dataset were considered (Figure 3A). However, Firmicutes ASVs showed higher community-wide diversity when host diversity was high (Figure 3B), while the Proteobacteria ASVs show less variation in diversity across locations (Figure 3C).

**Figure 1:**
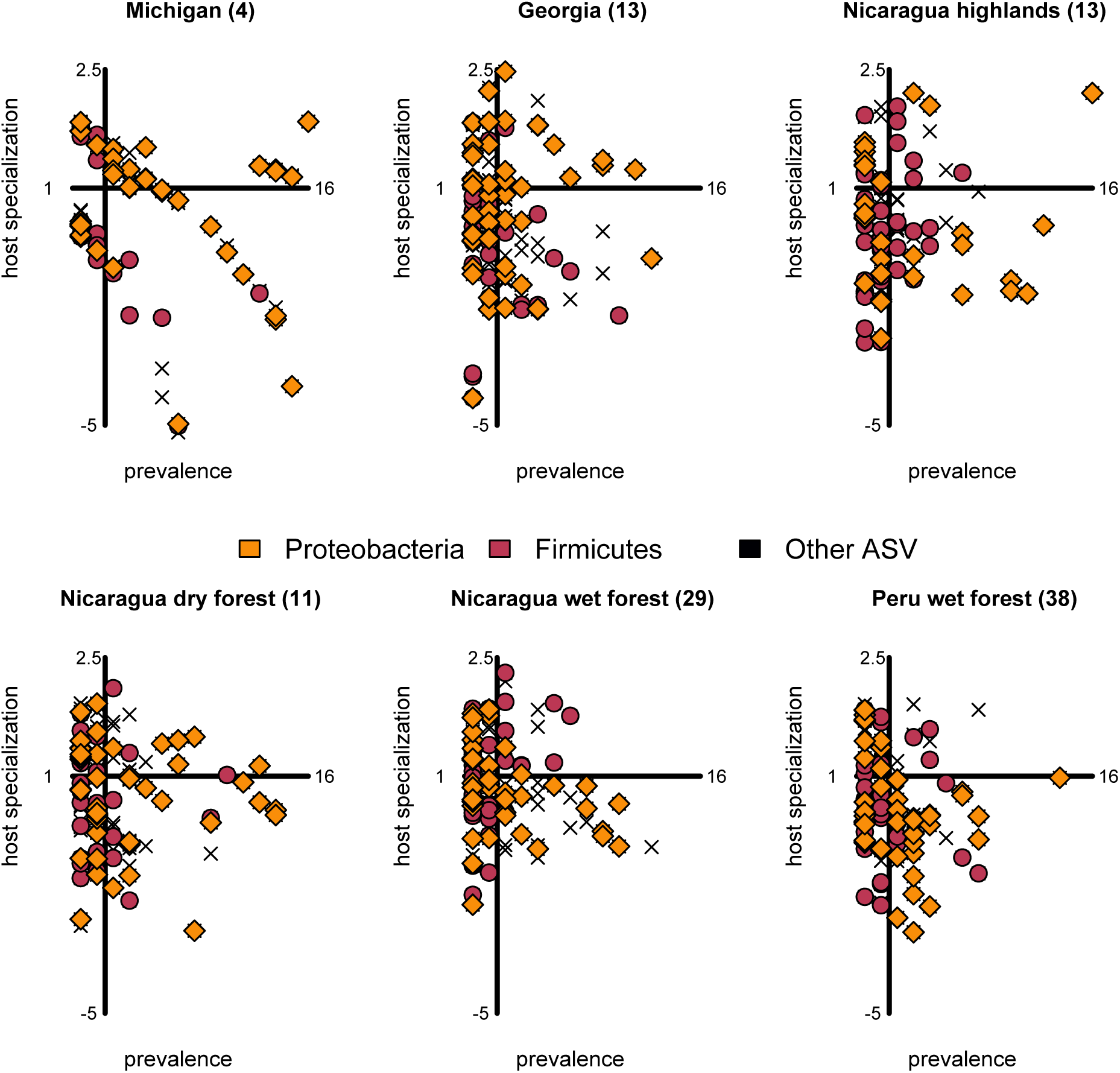
Abundance compared to the specialization index of ASVs across host communities. The x axis represents ASV abundance, while the y axis represents host specialization. The y axis is drawn at a value of x=2.5, the mean abundance value. The x axis is drawn at a specialization index of 0.

**Figure 2:**
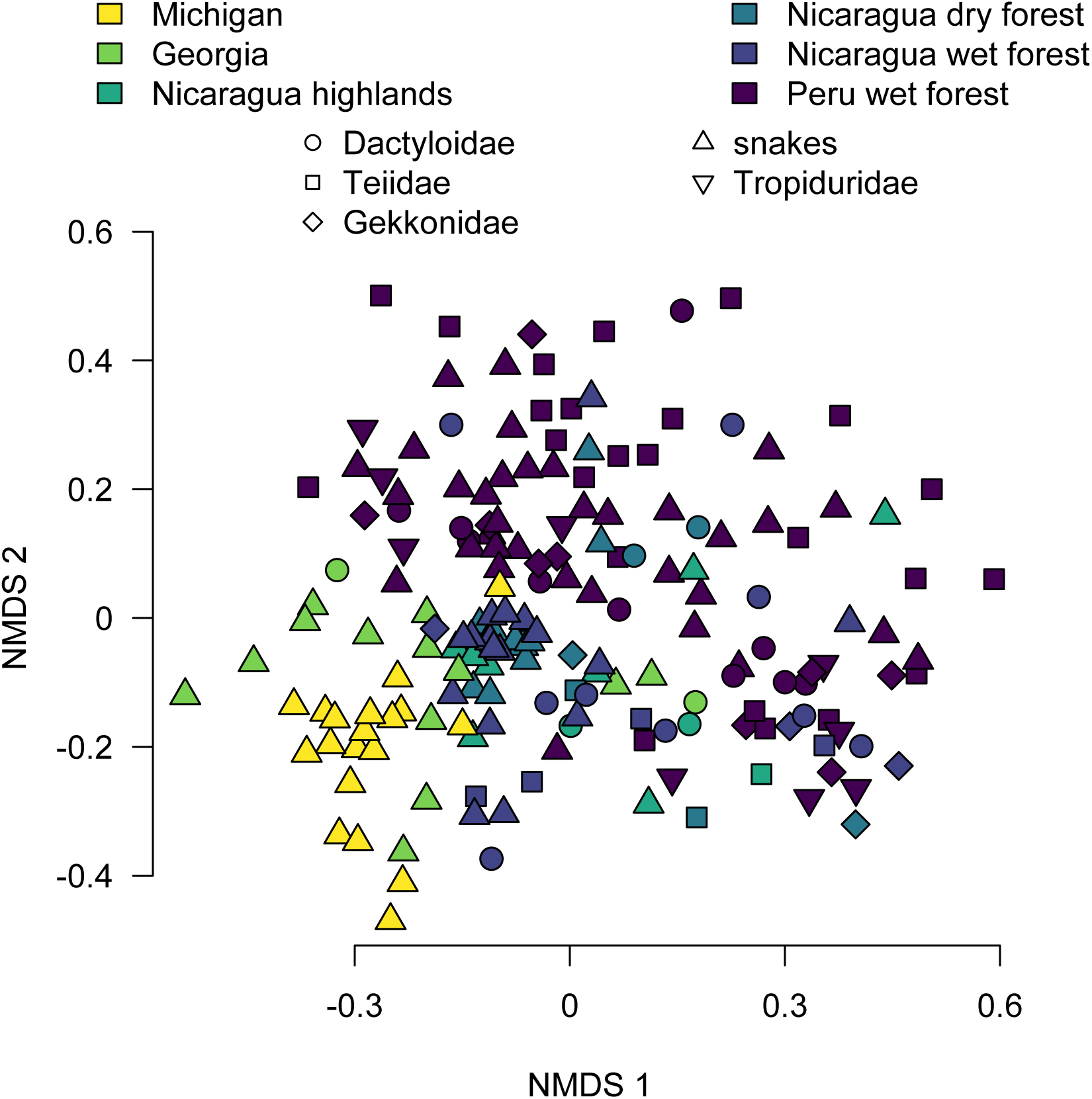
Pairwise Bray-Curtis distances between the microbiome communities of each host. Samples separate roughly by location, with some clustering by host phylogeny.

**Figure 3:**
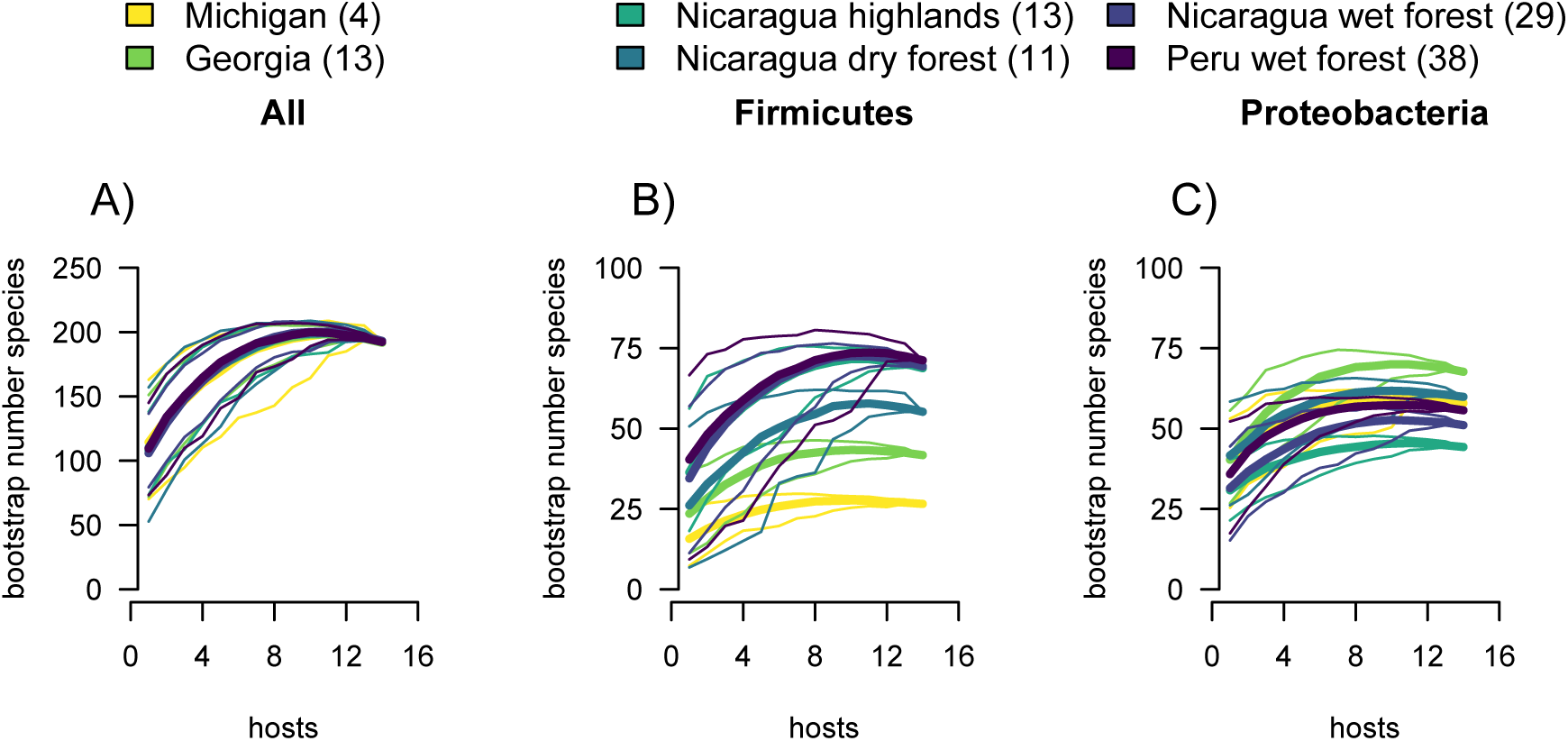
ASV lineage accumulation curves using jackknifing show similar predicted microbial diversity across equally sized samples of hosts in all locations (A). However, Firmicutes (the host-specific group) have higher community-wide diversity in high-host diversity communities (B), whereas the more generalist Proteobacteria show a different pattern not related to host diversity (C). Thick lines indicate mean values of the species accumulation curve, while thin lines show the 95% confidence interval of the jackknife values.

Total within-host bacterial diversity did not vary across locations. Shannon diversity values of individual microbiomes were not significantly different across sites when all ASVs were considered (p=0.89; Table S2 for per-location values of the metric). Firmicutes-only (p=0.0143) and Proteobacteria-only (p=0.0289) tests were also not significant at our Bonferroni cutoff. Incorporating phylogenetic information using the Faith’s diversity index amplified the differences between sites and bacterial phyla (Figure 4A; Table S2). The full communities again were not significantly different (p=0.181), while Firmicutes and Proteobacteria comparisons were significant (p=3.83×10^-16^; p < 2×10^-16^). Firmicutes diversity increased with host diversity, while Proteobacteria decreased (Figure 4A; Table S3 for Tukey test pairwise comparisons across sites). The ASV-specific host specialization index was significantly different between sites (p=3.19×10^-7^). Firmicutes ASV specialization (p=1.64×10^-6^) differed across locations, while Proteobacteria ASVs did not (p=0.285). This pattern was largely driven by higher specialization in the Nicaragua wet forest site, a possible sampling error (Figure 4B; Table S3). Overall, our results point toward specialist, low-prevalence strategies being more common in high host diversity communities, while lower-diversity host communities contain some extremely generalist, high prevalence lineages.

**Figure 4:**
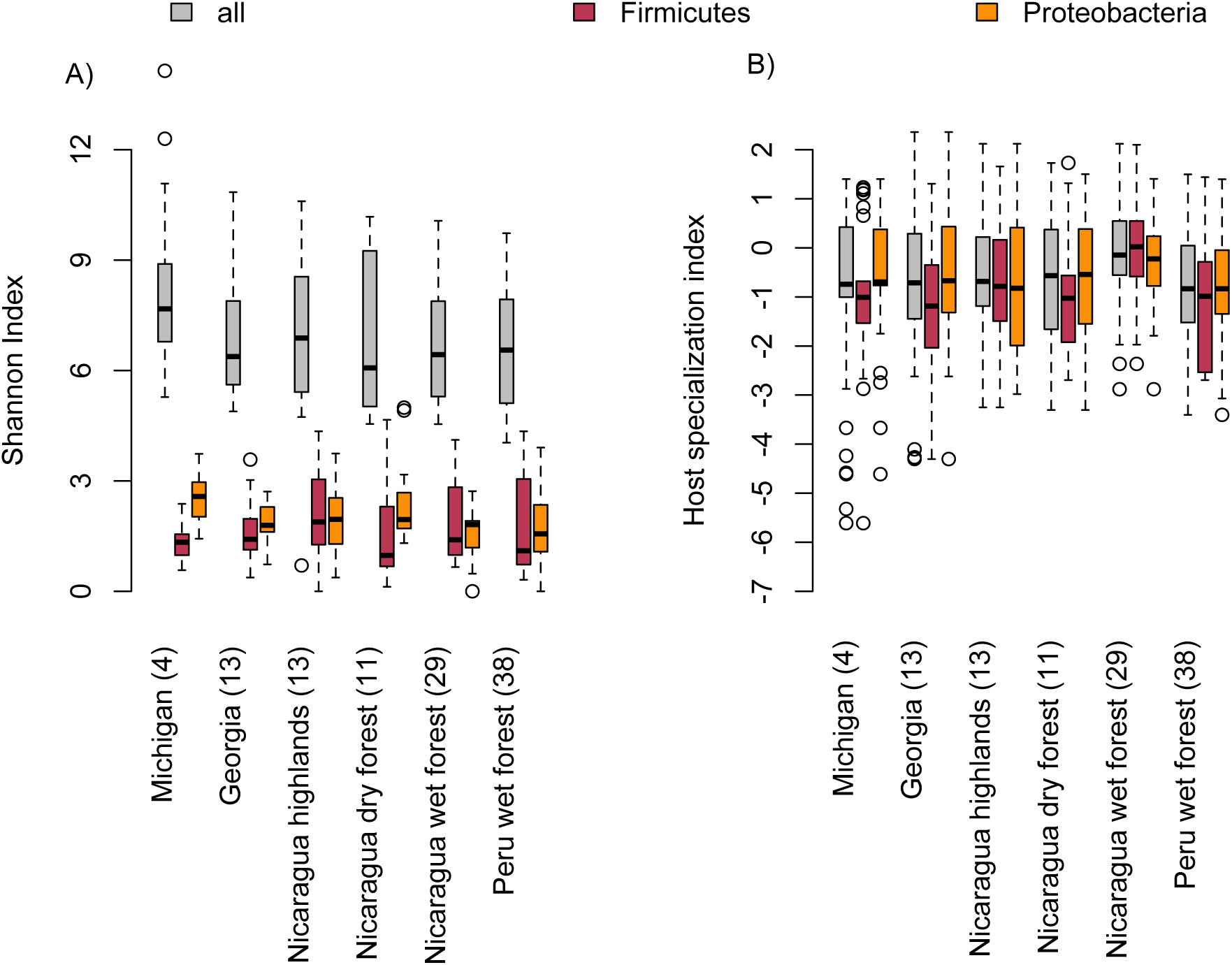
Microbiome diversity from low to high-diversity host populations (A) and ASV specialization (B). Full microbiome communities are not significantly different in host diversity across host communities using Faith’s phylogenetic diversity index (A). However, the diversity of microbes from a phylum enriched with host-specific taxa (Firmicutes) increases towards the tropics, while the diversity of generalist microbes (Proteobacteria) decreases. Accounting for the differences in host community diversity, the range of the host specialization index remains relatively consistent across sites (B).

### Within-community ASV co-occurrence patterns

To identify signatures of ecological interactions between ASVs, we found pairs of ASVs that cooccurred in hosts significantly more or less often than expected. We compared positively and negatively co-occurring pairs from 50 random, rarefied subsamples of the dataset, each with a paired null dataset. We found significantly more positively co-occurring pairs in high host diversity communities (ANOVA p < 2×10^-16^; Figure 5A; Table S3), and each location had significantly more pairs than the randomized dataset (Table S2 for t-test p-values). The number of negatively co-occurring pairs was also significantly different across host communities (ANOVA p < 2×10^-16^), but the pattern was less geographically ordered (Figure 5A; Table S3) and the real number of pairs was only significantly different from the randomized dataset in two locations (Georgia, p=1.678×10^-15^ and highland Nicaragua, p=2.134×10^-9^). The number of positively co-occurring pairs was significantly different from the number of negatively co-occurring pairs in every location (Table S2).

**Figure 5:**
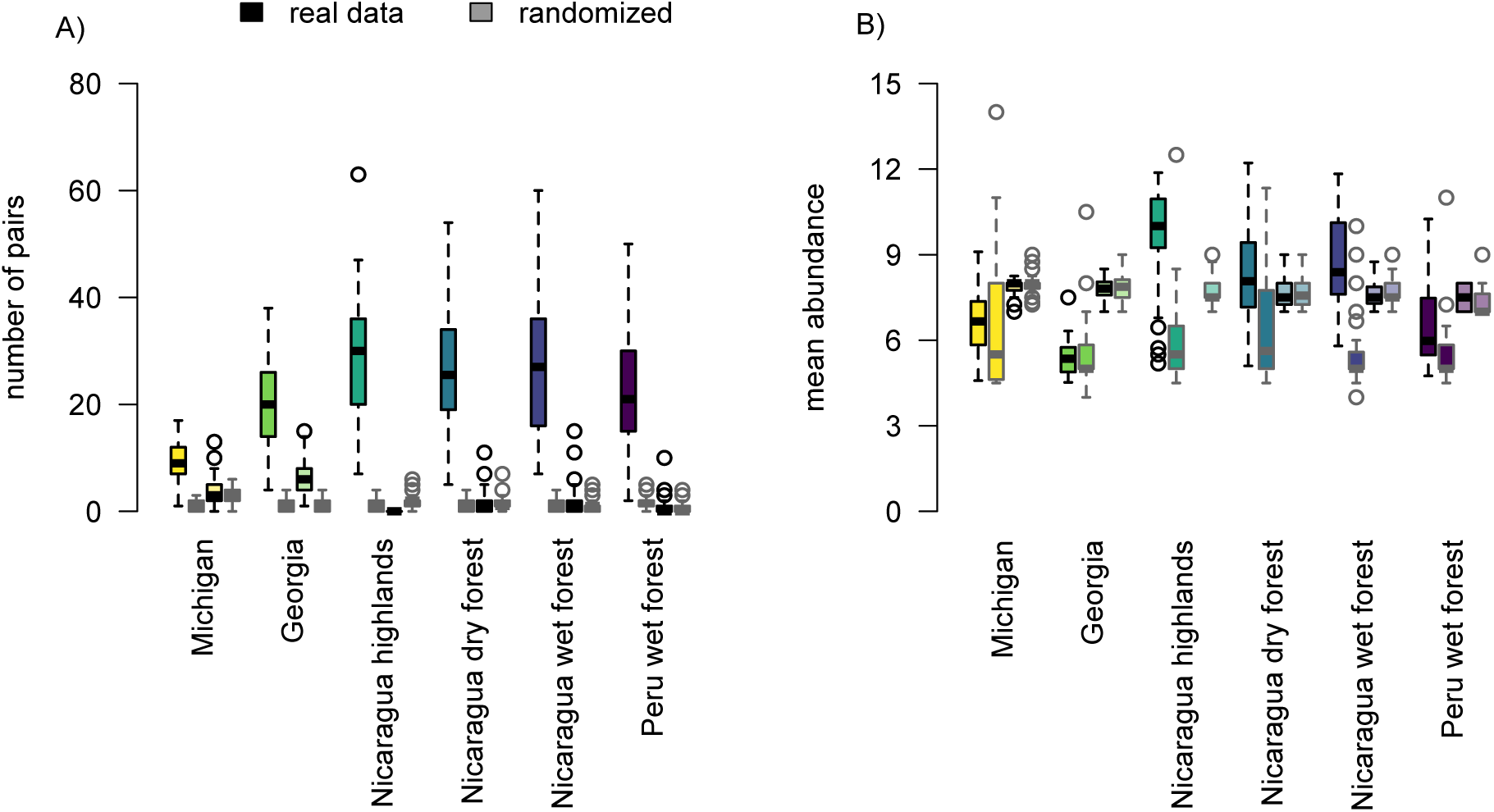
Interaction patterns across locations. Boxplots are colored by location. Black outlines indicate results from real data, grey lines show results from randomized data. Full-saturation colors (the first two boxes) show positive co-occurrence, while transparent colors (boxes three and four) show negative co-occurrence. (A) Pairs of positively co-occurring ASVs are more common in our observed data than the null dataset. Negatively co-occurring pairs are less common than positively co-occurring pairs and are similar to their frequency in the null dataset. Locations with higher host diversity have more positively co-occurring pairs. (B) ASVs in positively co-occurring pairs in the real dataset are more prevalent than those in the null dataset. Negatively co-occurring pairs have similar abundance to the null dataset.

Because the negatively co-occurring pairs were similar to the null dataset outcomes in most locations (Figure 5A), we reported results only for the positively co-occurring pairs in our further analyses. Positively co-occurring ASVs were more prevalent (occupied more hosts) in the real compared to the null datasets (Figure 5B; Table S2). Prevalence of the co-occurring ASVs was significantly different across locations (ANOVA p < 2×10^-16^), with Nicaraguan samples showing significantly higher prevalence than other locations (Figure 5B; Table S3).

We found that co-occurring ASV pairs were more phylogenetically similar in high-diversity host communities (ANOVA p=4.42×10^-14^). Comparison between sites showed significant differences between the Michigan and the Nicaraguan locations (Figure 6A; Table S3). The phylogenetic distances between co-occurring pairs were consistently larger than the mean nearest neighbor distances for each location, indicating that repeated within-host diversification was not a likely explanation for the pattern. Our low host diversity communities showed similar phylogenetic distances between pairs in the real and randomized datasets (Figure 6A). In all other locations, the real pairs were more phylogenetically similar than the pairs from the randomized datasets (Table S2). We found a significant excess of Proteobacteria-Proteobacteria pairs compared to the random expectation in every location except our lowest host diversity community (Figure 7; Table S2). Overall, our findings indicate that groups of co-occurring bacterial lineages may facilitate high prevalence and host generalism in high-diversity host communities.

**Figure 6:**
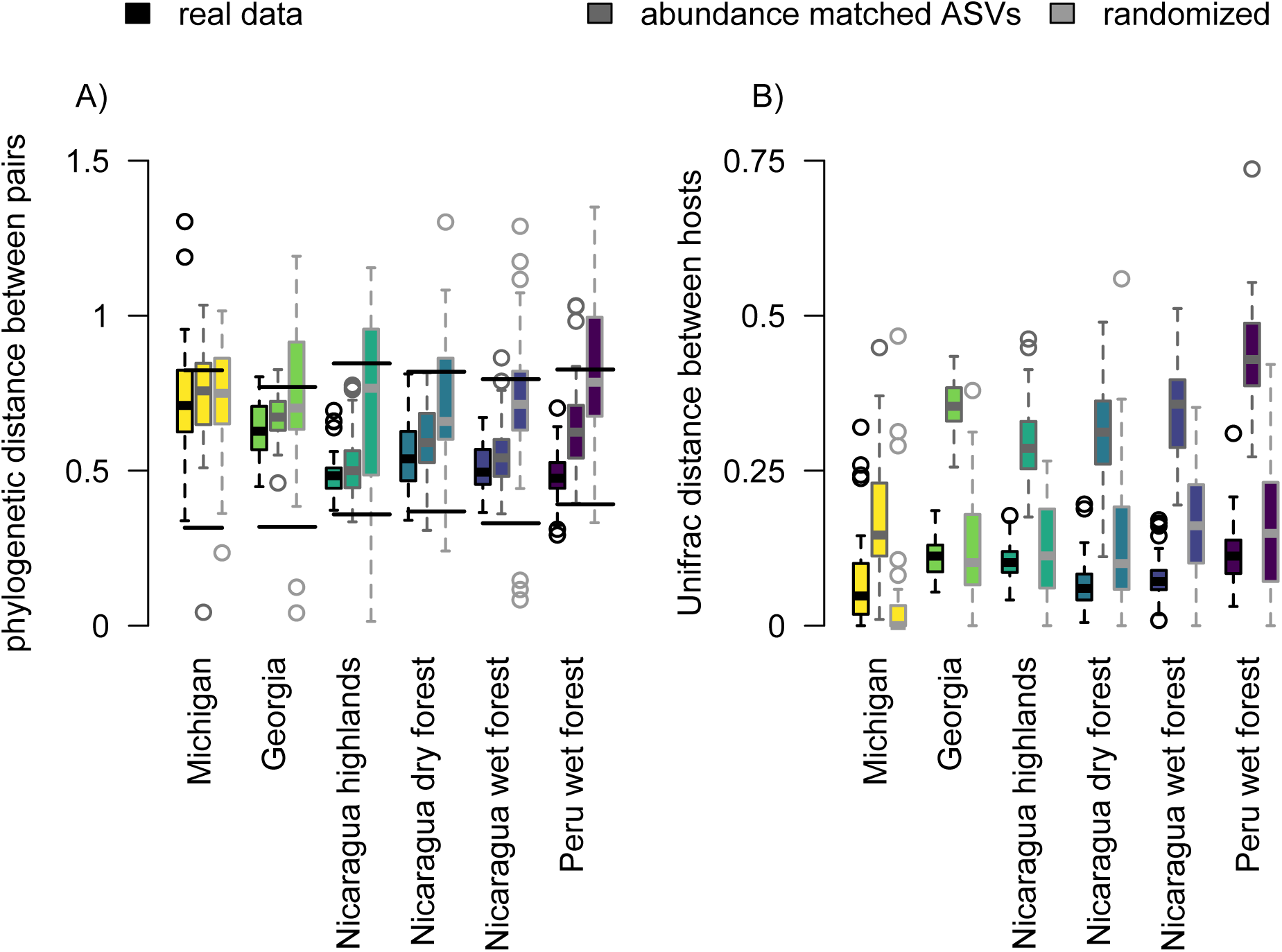
Characteristics of interacting ASVs. Boxes are colored by location. Black outlines are the real data from positively co-occurring pairs, medium grey outlines are data from non-interacting ASV pairs with the same prevalence as co-occurring pairs, and light grey outlines are positively co-occurring pairs from the null dataset. (A) Co-occurring pairs are more closely phylogenetically related than would be expected compared to the null dataset. This difference is partially explained for by accounting for prevalence, likely because the majority of high-prevalence ASVs are from the phylum Proteobacteria. This trend is more apparent in high host diversity communities. The lower set of horizontal lines indicate mean nearest neighbor distance for each location in a rarefied dataset. Since the mean distance between co-occurring pairs is greater than the nearest neighbor distance, within-host diversification is an unlikely explanation for our observed co-occurrence patterns. The upper set of lines represent the mean phylogenetic pairwise distance between ASVs in the rarefied dataset. (B) Host communities are more phylogenetically related than would be expected by chance (using Unifrac distance) compared to the hosts of the matched-abundance pairs. However, much of this variation is accounted for in the null dataset, since by definition co-occurring pairs share hosts. Notably, the host similarity in the real pairs does not increase in high-diversity host communities, while the host distances between matched-prevalence pairs increase with total host community diversity.

**Figure 7:**
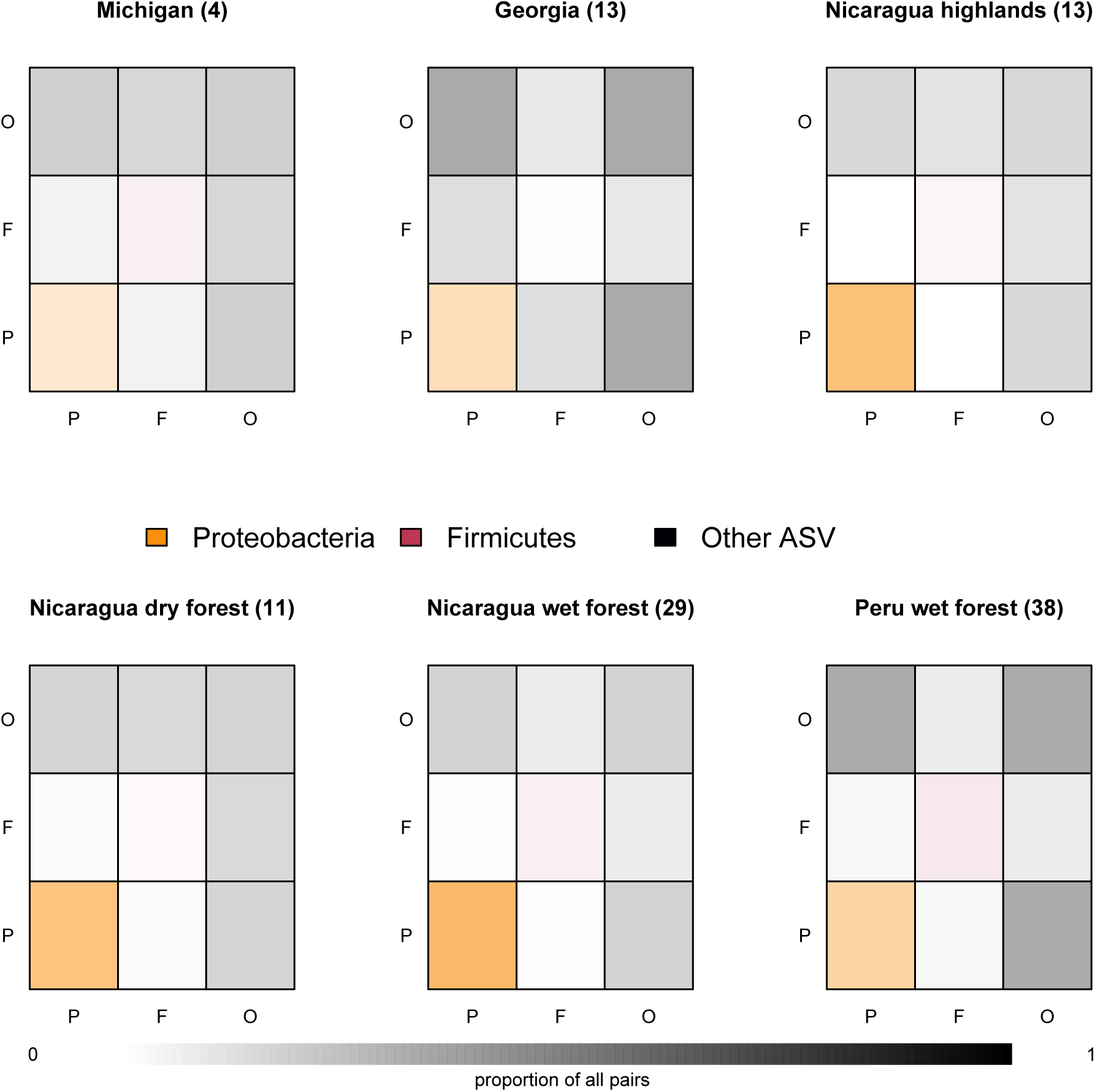
Proportion of co-occurring pairs made up of Proteobacteria, Firmicutes, and other phyla. Phyla are indicated by color of the squares and the letters (P, F, and O) on the margins. The proportion of each pair type is indicated by color intensity. In higher-diversity host communities, Proteobacteria-Proteobacteria pairs (orange, lower left) are predominant. Firmicutes-Firmicutes pairs (central square) are rare across all sites.

### Drivers of positive bacterial co-occurrence

When comparing the phylogenetic distance between co-occurring pairs of ASVs to prevalence-matched, randomly drawn pairs, only pairs from Peru were significantly different from the null dataset (Figure 6A; Table S2). To determine if overlap between co-occurring pairs could be driven by shared specialization on a host taxon, we tested host phylogenetic distance between pairs using the UniFrac distance metric. Host phylogenetic distance was significantly different across locations (ANOVA p=1.65×10^-6^), with a trend toward larger distances between hosts following overall host diversity (Figure 6B). No pairwise comparison was significant (Table S3). However, host diversity considering only co-occurring ASVs remained relatively consistent across locations, while the diversity of hosts in the prevalence-matched dataset rose in tandem with community host diversity. The real and null datasets were not significantly different with respect to host phylogenetic similarity in any location (Table S2), while the real hosts were significantly more phylogenetically similar relative to the hosts of the prevalence matched ASVs in all locations except Michigan (Table S2). This outcome is likely a feature of co-occurrence rather than host specialization, in that co-occurring ASVs by necessity share many hosts.

We found interacting modules of hosts and bacteria within our bipartite networks by identifying groups that share more links within a module than outside the module. Our community detection algorithm found that modularity increased significantly from low to high diversity host communities (ANOVA p < 2×10^-16^, Table S2, Table S3). Higher modularity indicated that ASVs were more specific to within-module hosts. Modularity of our randomized matrices also increased along this gradient, but the modularity of the randomized matrices was significantly lower than the modularity of the real matrices within each location (Table S2).

Across all locations, the real datasets had fewer, larger modules than the randomized datasets (Table S2). In the real data, both module number (p=1.83×10^-5^), number of ASVs per module (p=7.51×10^-6^), and number of hosts per module (p=2.41×10^-6^) differed significantly across locations, driven by high module numbers and low module size in our intermediate diversity, high elevation Nicaraguan site. These outcomes paralleled our finding of high numbers of co-occurring pairs of microbes to show the importance of bacteria-bacteria interactions in our dataset.

ASVs were more phylogenetically clustered in modules calculated from the real relative to the randomized datasets (Table S2). Our moderate and high host diversity communities had significantly greater phylogenetic clustering of ASVs relative to our low host diversity communities (ANOVA p < 2×10^-16^, Table S3 for pairwise comparisons). Hosts were significantly more similar in the real compared to randomized datasets across all sites. Host specialization values for ASVs within modules were significantly different across sites (ANOVA p < 2×10^-16^, Table S3 for pairwise comparisons), but in contrast to the ASV values, hosts were least phylogenetically clustered in our two most diverse communities. This pattern may reflect the importance of bacteria-bacteria interactions in overcoming limitations on generalism in high host diversity communities.

## Discussion

While some bacterial lineages demonstrate a remarkable ability to survive in many habitats, our analysis demonstrates a limit to that ability in this host-associated microbial system. In our two most diverse host communities, our most generalist ASVs showed lower prevalence and lower relative host generalism compared to the most generalist ASVs in less-diverse host communities (Figure 1; Figure 2). Our two dominant bacterial phyla, Firmicutes and Proteobacteria, had distinct patterns of host specialization. Proteobacteria were more generalist and achieved higher prevalence across both host individuals and species, while Firmicutes showed stronger specificity to host lineages (Gripp et al. 2011; Dandekar et al. 2012; Touchon et al. 2020). Firmicutes lineages have successfully established vertical transmission in other host taxa (Moeller et al. 2016, 2018). If they do so in our sampled communities, their transmission mode may facilitate specialization as a successful strategy in high host diversity communities (Figure 1). We identified higher-than-expected levels of positive co-occurrence between microbial lineages across all locations, but particularly in high host diversity communities (Figure 5). The ASVs that participated in these patterns were disproportionately from the phylum Proteobacteria (Figure 7). For each location, the interacting ASVs were some of the most prevalent in the community (Figure 6). Together, we showed that host-associated microbes were restricted in their host generalism, particularly as total host diversity increased. However, because co-occurrence also increased for these taxa in high host diversity communities, we speculate that microbe-microbe interactions may help to facilitate the generalist strategy even in challenging ecological circumstances (Figure 5B, Figure 7).

Our hypothesis that bacterial diversity would reflect host diversity within a community was not upheld when we considered all ASVs (Figure 3A). Although the diversity of host-specific Firmicutes lineages did increase with host diversity (Figure 3B), both per-host (Figure 4) and community-wide (Figure 3C), Proteobacteria diversity was lower in high-diversity host communities. This difference is likely driven by the absence of highly prevalent and host generalist Proteobacteria ASVs in high diversity host communities (Figure 2, Table S2), contradicting our hypothesis that high host diversity would drive ASVs to become either highly specialist, or highly generalist. Our data show patterns that directly contrast with free-living bacteria assemblages, in which generalist microbes are more diverse at southern latitudes (Zuo et al. 2023). This contrast may point to a fundamental difference in host filtering compared to free-living bacterial community assembly. However, these systems, like ours (Figure 2), showed that generalist bacterial lineages are more geographically structured than specialists (Liao et al. 2016; Hu et al. 2019; Luo et al. 2019; Zuo et al. 2023).

Our findings may indicate a relative advantage for specialist microbes in high-diversity communities. However, the trend toward higher Firmicutes diversity is not visible between our lowest diversity and mid-diversity host communities, suggesting that limits to generalism occur only at very high host species diversity. This pattern could be accounted for if ASVs were restricted in their total phylogenetic breadth of host taxa. However, all sites except our most northerly location had similar values for the maximum host generalism by an ASV (Figure 1), suggesting that the restriction in generalism in tropical communities is not due to an absolute incompatibility between hosts and generalist ASVs. We therefore hypothesize that specialists are competitively dominant compared to generalists in high host diversity communities. In this case, competition could constrain the realized niche of generalists while their fundamental niche remains broad. Similar competition-dependent specialization has been observed in plant-microbe systems (Lang et al. 2017), and is frequently observed among species in macro-scale communities (Looney et al. 2018; Lounibos and Juliano 2018; Jaszczuk et al. 2023). Future work will be necessary to determine whether this competition occurs directly through resource use, indirectly through modulation of host immune responses, or through some other mechanism (Råberg et al. 2006; Halliday et al. 2018).

We found that the ASVs that were able to achieve relatively high prevalence and host generalism in high-diversity host communities were disproportionately associated across hosts with one or more other bacterial lineage (Figure 5). The interacting ASVs were most often both from the phylum Proteobacteria, but tended to be more distantly related within that phylum (Figure 6A; Figure 7). Biological explanations for the pattern of co-occurring lineages include processes like cooperative biofilm formation or cross-feeding, both of which could buffer bacteria against host immune defenses or competition from specialists (McNally et al. 2014). Alternatively, the outcome could reflect shared food resources, if hosts have similar diets (Muegge et al. 2011; Delsuc et al. 2014; Song et al. 2019), or specialization on the same host lineages. However, we expect the total diversity of potential food items to be higher in the tropics, reducing the likelihood of extensive shared specialization relative to temperate communities (Andrew and Hughes 2004; Piel 2018). The relatively lower incidence of Firmicutes-Firmicutes pairs could point to a tradeoff between maintaining host specificity and participating in mutualistic interactions with other bacterial taxa. In total, our data indicate that patterns of host specificity are correlated with consistent and significant differences in community interactions between bacterial phyla.

Our study has important methodological caveats. First, we sampled locations only in one season. Patterns of host-bacteria and bacteria-bacteria interactions could change profoundly through time. Second, extraction methods can impact the taxa recovered from cloacal swabs (Hoffbeck et al. 2023), meaning that our sequenced samples are likely not a complete picture of the cloacal microbiome. Finally, we cannot directly test for evidence of interactions between bacteria with metabarcoding data. Metagenomic and metatranscriptomic studies will be necessary to find the mechanistic basis of the ecological patterns we describe.

## Conclusions

We demonstrate that even the most generalist host-associated bacteria are constrained in their generalism in extremely high diversity host communities. These limitations could be imposed by host immune responses, nutrient availability, or competition from other members of the microbial community (Bolnick et al. 2014; Coyte et al. 2015; Hansen et al. 2015; Holmes et al. 2017; Verster and Borenstein 2018). We hypothesize that the drivers of our observed patterns are a combination of these factors, in which microbes that are somewhat maladapted to a host environment are at a competitive disadvantage and can be excluded by better-adapted specialists.

Bacteria tend to reduce their genome size as they specialize (Baumler and Fang 2013; Sheppard et al. 2018), while larger genomes facilitate generalism and resilience to environmental perturbations. To get around this tradeoff and reap the benefits of a large metabolic repertoire without increasing genome size, bacterial lineages can develop stable interactions with other bacteria. Our data shows a strong signature of the importance of such interactions in high host diversity communities. If it is possible for bacteria to disperse as a unit, or if all lineages involved in the interaction are common in the environment, this type of interaction may be more successful than single-lineage generalism (Ben-Jacob et al. 2016; Natan et al. 2022).

We found that the preponderance of positive co-occurrence interactions, particularly in high host diversity communities, were between members of the phylum Proteobacteria (Figure 7). This stands in contrast to the overall pattern of co-occurring ASVs being habitat specialists across non-host associated habitats (Ma et al. 2020). In our dataset, the interacting pairs were relatively distantly related within their phylum (Figure 6A). We hypothesize that this pattern hints at a ‘Goldilocks zone’ of evolutionary divergence for mutualistic interactions. Closely related pairs might be more likely to compete than cooperate, while distantly related pairs may not share enough metabolic pathways in common to successfully facilitate each other. This outcome is in contradiction to our hypothesis that interactions would be stronger between host specialist microbes. Instead, our observed patterns may reflect a tradeoff between specializing on host interactions (used by Firmicutes) compared to investing in interactions with other bacteria (used by Proteobacteria).

## Acknowledgements

We appreciate the Matthaei Botanical Garden for granting access to their property. Samples were collected under Michigan State Scientific Collecting Permits 01-14-2016, 02-23-2017, and 06-18-2018. We appreciate help with sample collection in Georgia from Christian Cox. Samples were collected under a Scientific Collecting Permit from the Georgia Department of Natural Resources to Christian L. Cox (licensee #1000545789). For the Nicaraguan samples, we want to thank property owners and managers Sandra Castrillo (Refugio Bartola), Ramón Jiménez and family (Las Brisas del Mogotón), and Israel Molina Díaz (Laguna Asososca and Momotombo).

We also thank Heyddy Calderón Palma, Indiana Montoya, and René Castellón from the Ministerio de Ambiente y Recursos Naturales (MARENA) for their help obtaining research permits. Samples were collected under research permits by the national authority Ministerio de Ambiente y Recursos Naturales (MARENA) No. DGPNB-IC-025-2018. For sample collection in Peru, we would like to thank Ciara M. Sánchez-Paredes, Valia Herrera, Óscar Huacarpuma Aguilar, Edgar Iglesias Antonio, Eliz Lennia, César Macahuache Díaz, Greg Pandelis, Dan Rabosky, Alison Rabosky, Imani Russell, Roy Santa Cruz Farfán, Niery Tafur Olortegui, Pascal Title, Erick Vargas Laura, and Randi Villarcorta Díaz. This research was conducted under permits from the Peruvian government agency SERFOR (Servicio Nacional Forestal y de Fauna Silvestre; permit numbers: 029-2016-SERFOR-DGGSPFFS, 405-2016-SERFORDGGSPFFS, 116-2017-SERFOR-DGGSPFFS). Laboratory work for this project was carried out in the University of Michigan Biodiversity Laboratory. This research was supported by startup funds from the University of Michigan to ADR, research funds from the Rackham Graduate School to JGL and IAH, funds from the University of Michigan Department of Ecology and Evolutionary Biology to IAH, and from the Packard Foundation to Dan Rabosky. National Institutes of Health and National Institute of Allergy and Infectious Diseases Award T32Il45821 funded IAH during analysis.

## Notes

### Competing Interest Statement

The authors have declared no competing interest.

### Summary of Updates

We have submitted a revised version to correct Figure 7, which lacked color when converted to a pdf.

## Works Cited

Aijuka, M., and E. M. Buys. 2019. Persistence of foodborne diarrheagenic Escherichia coli in the agricultural and food production environment: Implications for food safety and public health. Food Microbiology 82:363–370.

Ambrosini, R., M. Corti, A. Franzetti, M. Caprioli, D. Rubolini, V. M. Motta, A. Costanzo, N. Saino, and I. Gandolfi. 2019. Cloacal microbiomes and ecology of individual barn swallows. FEMS Microbiology Ecology 95:fiz061.

Andrew, N. R., and L. Hughes. 2004. Species diversity and structure of phytophagous beetle assemblages along a latitudinal gradient: predicting the potential impacts of climate change. Ecological Entomology 29:527–542.

Baumler, A., and F. C. Fang. 2013. Host Specificity of Bacterial Pathogens. Cold Spring Harbor Perspectives in Medicine 3:a010041–a010041.

Beckett, S. J. 2016. Improved community detection in weighted bipartite networks. R. Soc. open sci. 3:140536.

Bell, T. H., and T. Bell. 2020. Many roads to bacterial generalism. FEMS Microbiology Ecology 97:fiaa240.

Ben-Jacob, E., A. Finkelshtein, G. Ariel, and C. Ingham. 2016. Multispecies Swarms of Social Microorganisms as Moving Ecosystems. Trends in Microbiology 24:257–269.

Bisanz, J. E. 2018. qiime2R: Importing QIIME2 artifacts and associated data into R sessions. Bodawatta, K. H., B. Koane, G. Maiah, K. Sam, M. Poulsen, and K. A. Jønsson. 2021. Species-specific but not phylosymbiotic gut microbiomes of New Guinean passerine birds are shaped by diet and flight-associated gut modifications. Proc. R. Soc. B. 288:rspb.2021.0446, 20210446.

Bolnick, D. I., L. K. Snowberg, J. G. Caporaso, C. Lauber, R. Knight, and W. E. Stutz. 2014. Major H istocompatibility C omplex class II b polymorphism influences gut microbiota composition and diversity. Mol Ecol 23:4831–4845.

Bolyen, E., J. R. Rideout, M. R. Dillon, N. A. Bokulich, C. C. Abnet, G. A. Al-Ghalith, H. Alexander, E. J. Alm, M. Arumugam, F. Asnicar, Y. Bai, J. E. Bisanz, K. Bittinger, A. Brejnrod, C. J. Brislawn, C. T. Brown, B. J. Callahan, A. M. Caraballo-Rodríguez, J. Chase, E. K. Cope, R. Da Silva, C. Diener, P. C. Dorrestein, G. M. Douglas, D. M. Durall, C. Duvallet, C. F. Edwardson, M. Ernst, M. Estaki, J. Fouquier, J. M. Gauglitz, S. M. Gibbons, D. L. Gibson, A. Gonzalez, K. Gorlick, J. Guo, B. Hillmann, S. Holmes, H. Holste, C. Huttenhower, G. A. Huttley, S. Janssen, A. K. Jarmusch, L. Jiang, B. D. Kaehler, K. B. Kang, C. R. Keefe, P. Keim, S. T. Kelley, D. Knights, I. Koester, T. Kosciolek, J. Kreps, M. G. I. Langille, J. Lee, R. Ley, Y.-X. Liu, E. Loftfield, C. Lozupone, M. Maher, C. Marotz, B. D. Martin, D. McDonald, L. J. McIver, A. V. Melnik, J. L. Metcalf, S. C. Morgan, J. T. Morton, A. T. Naimey, J. A. Navas-Molina, L. F. Nothias, S. B. Orchanian, T. Pearson, S. L. Peoples, D. Petras, M. L. Preuss, E. Pruesse, L. B. Rasmussen, A. Rivers, M. S. Robeson, P. Rosenthal, N. Segata, M. Shaffer, A. Shiffer, R. Sinha, S. J. Song, J. R. Spear, A. D. Swafford, L. R. Thompson, P. J. Torres, P. Trinh, A. Tripathi, P. J. Turnbaugh, S. Ul-Hasan, J. J. J. Van Der Hooft, F. Vargas, Y. Vázquez-Baeza, E. Vogtmann, M. Von Hippel, W. Walters, Y. Wan, M. Wang, J. Warren, K. C. Weber, C. H. D. Williamson, A. D. Willis, Z. Z. Xu, J. R. Zaneveld, Y. Zhang, Q. Zhu, R. Knight, and J. G. Caporaso. 2019. Reproducible, interactive, scalable and extensible microbiome data science using QIIME 2. Nat Biotechnol 37:852–857.

Bono, L. M., L. B. Smith, D. W. Pfennig, and C. L. Burch. 2017. The emergence of performance trade-offs during local adaptation: insights from experimental evolution. Molecular Ecology 26:1720–1733.

Bordes, F., J. F. Guégan, and S. Morand. 2011. Microparasite species richness in rodents is higher at lower latitudes and is associated with reduced litter size. Oikos 120:1889–1896.

Bunker, M. E., G. Elliott, H. Heyer-Gray, M. O. Martin, A. E. Arnold, and S. L. Weiss. 2021. Vertically transmitted microbiome protects eggs from fungal infection and egg failure. anim microbiome 3:43.

Bunker, M. E., and S. L. Weiss. 2024. The reproductive microbiome and maternal transmission of microbiota via eggs in *Sceloporus virgatus*. FEMS Microbiology Ecology 100:fiae011.

Callahan, B. J., P. J. McMurdie, M. J. Rosen, A. W. Han, A. J. A. Johnson, and S. P. Holmes. 2016. DADA2: High-resolution sample inference from Illumina amplicon data. Nat Methods 13:581–583.

Chen, J., X. Zhang, L. Yang, and L. Zhang. 2023. GUniFrac: Generalized UniFrac Distances Distance-Based Multivariate Methods and Feature-Based Univariate Methods for Microbiome Data Analysis.

Chen, Y.-J., P. M. Leung, J. L. Wood, S. K. Bay, P. Hugenholtz, A. J. Kessler, G. Shelley, D. W. Waite, A. E. Franks, P. L. M. Cook, and C. Greening. 2021. Metabolic flexibility allows bacterial habitat generalists to become dominant in a frequently disturbed ecosystem. The ISME Journal 15:2986–3004.

Clark, N. J. 2018. Phylogenetic uniqueness, not latitude, explains the diversity of avian blood parasite communities worldwide. Global Ecol Biogeogr 27:744–755.

Cobian, G. M., C. P. Egan, and A. S. Amend. 2019. Plant–microbe specificity varies as a function of elevation. The ISME Journal 13:2778–2788.

Colston, T. J. 2017. Gut microbiome transmission in lizards. Molecular Ecology 26:972–974.

Colston, T. J., and C. R. Jackson. 2016. Microbiome evolution along divergent branches of the vertebrate tree of life: what is known and unknown. Molecular Ecology 25:3776–3800.

Colston, T. J., B. P. Noonan, and C. R. Jackson. 2015. Phylogenetic analysis of bacterial communities in different regions of the gastrointestinal tract of *Agkistrodon piscivorus*, the cottonmouth snake. PLOS ONE 10:e0128793.

Coyte, K. Z., J. Schluter, and K. R. Foster. 2015. The ecology of the microbiome: Networks, competition, and stability. Science 350:663–666.

Curtiss, J. B., A. M. Leone, J. F. X. Wellehan, J. A. Emerson, E. W. Howerth, and L. L. Farina. 2015. Renal and cloacal cryptosporidiosis ( *Cryptosporidium* avian genotype V) in a Major Mitchell’s cockatoo ( *Lophochroa leadbeateri* ). Journal of Zoo and Wildlife Medicine 46:934–937.

Dandekar, T., F. Astrid, P. Jasmin, and M. Hensel. 2012. Salmonella enterica: a surprisingly well-adapted intracellular lifestyle. Front. Microbio. 3.

Delsuc, F., J. L. Metcalf, L. Wegener Parfrey, S. J. Song, A. González, and R. Knight. 2014. Convergence of gut microbiomes in myrmecophagous mammals. Molecular Ecology 23:1301–1317.

DeSantis, T. Z., P. Hugenholtz, N. Larsen, M. Rojas, E. L. Brodie, K. Keller, T. Huber, D. Dalevi, P. Hu, and G. L. Andersen. 2006. Greengenes, a Chimera-Checked 16S rRNA Gene Database and Workbench Compatible with ARB. Appl Environ Microbiol 72:5069–5072.

Dormann, C. F., and R. Strauss. 2014. A method for detecting modules in quantitative bipartite networks. Methods Ecol Evol 5:90–98.

Dormann, C., J. Fruend, N. Bleuthgen, and B. Gruber. 2008. Introducing the bipartite Package: Analysing Ecological Networks. R News 8/2:8–11.

Eren, A. M., M. L. Sogin, H. G. Morrison, J. H. Vineis, J. C. Fisher, R. J. Newton, and S. L. McLellan. 2015. A single genus in the gut microbiome reflects host preference and specificity. The ISME Journal 9:90–100.

Griffith, D. M., J. A. Veech, and C. J. Marsh. 2016. **cooccur** : Probabilistic Species Co-Occurrence Analysis in *R*. J. Stat. Soft. 69.

Gripp, E., D. Hlahla, X. Didelot, F. Kops, S. Maurischat, K. Tedin, T. Alter, L. Ellerbroek, K. Schreiber, D. Schomburg, T. Janssen, P. Bartholomäus, D. Hofreuter, S. Woltemate, M. Uhr, B. Brenneke, P. Grüning, G. Gerlach, L. Wieler, S. Suerbaum, and C. Josenhans. 2011. Closely related Campylobacter jejuni strains from different sources reveal a generalist rather than a specialist lifestyle. BMC Genomics 12:584.

Grond, K., B. K. Sandercock, A. Jumpponen, and L. H. Zeglin. 2018. The avian gut microbiota: community, physiology and function in wild birds. J Avian Biol e01788.

Halliday, F. W., J. Umbanhowar, and C. E. Mitchell. 2018. A host immune hormone modifies parasite species interactions and epidemics: insights from a field manipulation. Proc. R. Soc. B. 285:20182075.

Hansen, N. C. K., E. Avershina, L. T. Mydland, J. A. Næsset, D. Austbø, B. Moen, I. Måge, and K. Rudi. 2015. High nutrient availability reduces the diversity and stability of the equine caecal microbiota. Microbial Ecology in Health & Disease 26.

Hernandez, J., C. Hucul, E. Reasor, T. Smith, J. W. McGlothlin, D. C. Haak, L. K. Belden, and I. T. Moore. 2021. Assessing age, breeding stage, and mating activity as drivers of variation in the reproductive microbiome of female tree swallows. Ecology and Evolution 11:11398– 11413.

Hoffbeck, C., D. M. R. L. Middleton, N. J. Nelson, and M. W. Taylor. 2023. 16S RRNA gene-based meta-analysis of the reptile gut microbiota reveals environmental effects, host influences and a limited core microbiota. Molecular Ecology 32:6044–6058.

Holmes, A. J., Y. V. Chew, F. Colakoglu, J. B. Cliff, E. Klaassens, M. N. Read, S. M. Solon-Biet, A. C. McMahon, V. C. Cogger, K. Ruohonen, D. Raubenheimer, D. G. Le Couteur, and S. J. Simpson. 2017. Diet-Microbiome Interactions in Health Are Controlled by Intestinal Nitrogen Source Constraints. Cell Metabolism 25:140–151.

Hu, A., H. Wang, M. Cao, A. Rashid, M. Li, and C.-P. Yu. 2019. Environmental Filtering Drives the Assembly of Habitat Generalists and Specialists in the Coastal Sand Microbial Communities of Southern China. Microorganisms 7:598.

IUCN. 2023. The IUCN Red List of Threatened Species.

Jaszczuk, I., W. Kotowski, Ł. Kozub, J. Kreyling, and E. Jabłońska. 2023. Physiological responses of fen mosses along a nitrogen gradient point to competition restricting their fundamental niches. Oikos 2023:e09336.

Johne, R., A. Konrath, M.-E. Krautwald-Junghanns, E. F. Kaleta, H. Gerlach, and H. Müller. 2002. Herpesviral, but no papovaviral sequences, are detected in cloacal papillomas of parrots. Archives of Virology 147:1869–1880.

Kembel, S. W., P. D. Cowan, M. R. Helmus, W. K. Cornwell, H. Morlon, D. D. Ackerly, S. P. Blomberg, and C. O. Webb. 2010. Picante: R tools for integrating phylogenies and ecology. Bioinformatics 26:1463–1464.

Kohl, K. D., A. Brun, M. Magallanes, J. Brinkerhoff, A. Laspiur, J. C. Acosta, E. Caviedes-Vidal, and S. R. Bordenstein. 2017. Gut microbial ecology of lizards: insights into diversity in the wild, effects of captivity, variation across gut regions and transmission. Mol Ecol 26:1175–1189.

Kozich, J. J., S. L. Westcott, N. T. Baxter, S. K. Highlander, and P. D. Schloss. 2013. Development of a dual-Index sequencing strategy and curation pipeline for analyzing amplicon sequence data on the MiSeq Illumina sequencing platform. Applied and Environmental Microbiology 79:5112–5120.

Lang, J., A. Vigouroux, A. El Sahili, A. Kwasiborski, M. Aumont-Nicaise, Y. Dessaux, J. A. Shykoff, S. Moréra, and D. Faure. 2017. Fitness costs restrict niche expansion by generalist niche-constructing pathogens. The ISME Journal 11:374–385.

Lawrence, E. R., and D. J. Fraser. 2020. Latitudinal biodiversity gradients at three levels: Linking species richness, population richness and genetic diversity. Global Ecol Biogeogr 29:770– 788.

Lettink, M., and K. M. Hare. 2016. Sampling techniques for New Zealand lizards. Pp. 269–291 in New Zealand Lizards. Springer.

Ley, R. E., M. Hamady, C. Lozupone, P. J. Turnbaugh, R. R. Ramey, J. S. Bircher, M. L. Schlegel, T. A. Tucker, M. D. Schrenzel, R. Knight, and J. I. Gordon. 2008. Evolution of mammals and their gut microbes. Science 320:1647–1651.

Li, T., Y. Yang, H. Li, and C. Li. 2022. Mixed-Mode Bacterial Transmission via Eggshells in an Oviparous Reptile Without Parental Care. Front. Microbiol. 13:911416.

Liao, J., X. Cao, L. Zhao, J. Wang, Z. Gao, M. C. Wang, and Y. Huang. 2016. The importance of neutral and niche processes for bacterial community assembly differs between habitat generalists and specialists. FEMS Microbiology Ecology 92:fiw174.

Liu, X., and T. Murata. 2010. An Efficient Algorithm for Optimizing Bipartite Modularity in Bipartite Networks. Journal of Advanced Computational Intelligence and Intelligent Informatics (JACIII) 14.

Looney, C. E., A. W. D’Amato, S. Fraver, B. J. Palik, and L. E. Frelich. 2018. Interspecific competition limits the realized niche of *Fraxinus nigra* along a waterlogging gradient. Can. J. For. Res. 48:1292–1301.

Lounibos, L. P., and S. A. Juliano. 2018. Where vectors collide: the importance of mechanisms shaping the realized niche for modeling ranges of invasive Aedes mosquitoes. Biol Invasions 20:1913–1929.

Lozupone, C., and R. Knight. 2005. UniFrac: a New Phylogenetic Method for Comparing Microbial Communities. Appl Environ Microbiol 71:8228–8235.

Luo, Z., J. Liu, P. Zhao, T. Jia, C. Li, and B. Chai. 2019. Biogeographic Patterns and Assembly Mechanisms of Bacterial Communities Differ Between Habitat Generalists and Specialists Across Elevational Gradients. Front. Microbiol. 10:169.

Ma, B., Y. Wang, S. Ye, S. Liu, E. Stirling, J. A. Gilbert, K. Faust, R. Knight, J. K. Jansson, C. Cardona, L. Röttjers, and J. Xu. 2020. Earth microbial co-occurrence network reveals interconnection pattern across microbiomes. Microbiome 8:82.

Mallott, E. K., and K. R. Amato. 2021. Host specificity of the gut microbiome. Nat Rev Microbiol 19:639–653.

Mariadassou, M., S. Pichon, and D. Ebert. 2015. Microbial ecosystems are dominated by specialist taxa. Ecology Letters 18:974–982.

Maritan, E., A. Quagliariello, E. Frago, T. Patarnello, and M. E. Martino. 2024. The role of animal hosts in shaping gut microbiome variation. Phil. Trans. R. Soc. B 379:20230071.

Martínez-Fonseca, J. G., I. A. Holmes, J. Sunyer, E. P. Westeen, M. R. Grundler, P. A. Cerda, M. A. Fernández-Mena, J. C. Loza-Molina, I. V. Monagan Jr., D. Nondorf, G. G. Pandelis, and A. R. Davis Rabosky. 2024. A collection and analysis of amphibians and reptiles from Nicaragua with new country and departmental records. CheckList 20:58–125.

McNally, L., M. Viana, and S. P. Brown. 2014. Cooperative secretions facilitate host range expansion in bacteria. Nat Commun 5:4594.

Moeller, A. H., A. Caro-Quintero, D. Mjungu, A. V. Georgiev, E. V. Lonsdorf, M. N. Muller, A. E. Pusey, M. Peeters, B. H. Hahn, and H. Ochman. 2016. Cospeciation of gut microbiota with hominids. Science 353:380–382.

Moeller, A. H., T. A. Suzuki, M. Phifer-Rixey, and M. W. Nachman. 2018. Transmission modes of the mammalian gut microbiota. Science 362:453–457.

Mouftah, S. F., J. F. Cobo-Díaz, A. Álvarez-Ordóñez, A. Mousa, J. K. Calland, B. Pascoe, S. K. Sheppard, and M. Elhadidy. 2021. Stress resistance associated with multi-host transmission and enhanced biofilm formation at 42 °C among hyper-aerotolerant generalist Campylobacter jejuni. Food Microbiology 95:103706.

Muegge, B. D., J. Kuczynski, D. Knights, J. C. Clemente, A. González, L. Fontana, B. Henrissat, R. Knight, and J. I. Gordon. 2011. Diet Drives Convergence in Gut Microbiome Functions Across Mammalian Phylogeny and Within Humans. Science 332:970–974.

Natan, G., V. M. Worlitzer, G. Ariel, and A. Be’er. 2022. Mixed-species bacterial swarms show an interplay of mixing and segregation across scales. Sci Rep 12:16500.

Oksanen, J., F. Blanchet, M. Friendly, R. Kindt, P. Legendre, D. McGlinn, P. Minchin, R. O’Hara, G. Simpson, P. Solymos, M. Stevens, E. Szocs, and H. Wagner. 2018. vegan: Community Ecology Package.

Pebesma, E. 2018. Simple Features for R: Standardized Support for Spatial Vector Data. The R Journal 10:439–446.

Pebesma, E., and R. Bivand. 2023. Spatial Data Science: With Applications in R. Chapman and Hall/CRC.

Piel, W. H. 2018. The global latitudinal diversity gradient pattern in spiders. Journal of Biogeography 45:1896–1904.

Procheş, Ş., J. R. U. Wilson, and R. M. Cowling. 2006. How much evolutionary history in a 10×10 m plot? Proc. R. Soc. B. 273:1143–1148.

Råberg, L., J. C. De Roode, A. S. Bell, P. Stamou, D. Gray, and A. F. Read. 2006. The Role of Immune-Mediated Apparent Competition in Genetically Diverse Malaria Infections. The American Naturalist 168:41–53.

Sanders, J. G., D. D. Sprockett, Y. Li, D. Mjungu, E. V. Lonsdorf, J.-B. N. Ndjango, A. V. Georgiev, J. A. Hart, C. M. Sanz, D. B. Morgan, M. Peeters, B. H. Hahn, and A. H. Moeller. 2023. Widespread extinctions of co-diversified primate gut bacterial symbionts from humans. Nat Microbiol 8:1039–1050.

Sant, D. G., L. C. Woods, J. J. Barr, and M. J. McDonald. 2021. Host diversity slows bacteriophage adaptation by selecting generalists over specialists. Nat Ecol Evol 5:350–359.

Scott, J. J., T. C. Adam, A. Duran, D. E. Burkepile, and D. B. Rasher. 2020. Intestinal microbes: an axis of functional diversity among large marine consumers. Proc. R. Soc. B. 287:20192367.

Sheppard, S. K., D. S. Guttman, and J. R. Fitzgerald. 2018. Population genomics of bacterial host adaptation. Nat Rev Genet 19:549–565.

Smith, E. P., and G. Van Belle. 1984. Nonparametric Estimation of Species Richness. Biometrics 40:119.

Song, S. J., J. G. Sanders, D. T. Baldassarre, J. A. Chaves, N. S. Johnson, A. J. Piaggio, M. J. Stuckey, E. Nováková, J. L. Metcalf, B. B. Chomel, A. Aguilar-Setién, R. Knight, and V. J. McKenzie. 2019. Is there convergence of gut microbes in blood-feeding vertebrates? Phil. Trans. R. Soc. B 374:20180249.

Song, S. J., J. G. Sanders, F. Delsuc, J. Metcalf, K. Amato, M. W. Taylor, F. Mazel, H. L. Lutz, K. Winker, G. R. Graves, G. Humphrey, J. A. Gilbert, S. J. Hackett, K. P. White, H. R. Skeen, S. M. Kurtis, J. Withrow, T. Braile, M. Miller, K. G. McCracken, J. M. Maley, V. O. Ezenwa, A. Williams, J. M. Blanton, V. J. McKenzie, and R. Knight. 2020. Comparative analyses of vertebrate gut microbiomes reveal convergence between birds and bats. mBio 11:e02901–19.

Sriswasdi, S., C. Yang, and W. Iwasaki. 2017. Generalist species drive microbial dispersion and evolution. Nat Commun 8:1162.

Styles, D. K., E. K. Tomaszewski, L. A. Jaeger, and D. N. Phalen. 2004. Psittacid herpesviruses associated with mucosal papillomas in neotropical parrots. Virology 325:24–35.

Tillis, S. B., M. E. Iredale, A. L. Childress, E. A. Graham, J. F. X. Wellehan, R. Isaza, and R. J. Ossiboff. 2021. Oral, cloacal, and hemipenal actinomycosis in captive ball pythons (*Python regius*). Front. Vet. Sci. 7:594600.

Touchon, M., A. Perrin, J. A. M. De Sousa, B. Vangchhia, S. Burn, C. L. O’Brien, E. Denamur, D. Gordon, and E. P. Rocha. 2020. Phylogenetic background and habitat drive the genetic diversification of Escherichia coli. PLoS Genet 16:e1008866.

Van Der Valk, T., F. Vezzi, M. Ormestad, L. Dalén, and K. Guschanski. 2020. Index hopping on the Illumina HiseqX platform and its consequences for ancient DNA studies. Molecular Ecology Resources 20:1171–1181.

Van Veelen, H. P. J., J. Falcão Salles, K. D. Matson, M. Van Der Velde, and B. I. Tieleman. 2020. Microbial environment shapes immune function and cloacal microbiota dynamics in zebra finches Taeniopygia guttata. anim microbiome 2:21.

Van Veelen, H. P. J., J. Falcao Salles, and B. I. Tieleman. 2017. Multi-level comparisons of cloacal, skin, feather and nest-associated microbiota suggest considerable influence of horizontal acquisition on the microbiota assembly of sympatric woodlarks and skylarks. Microbiome 5:156.

Veech, J. A. 2013. A probabilistic model for analysing species co-occurrence. Global Ecology and Biogeography 22:252–260.

Verster, A. J., and E. Borenstein. 2018. Competitive lottery-based assembly of selected clades in the human gut microbiome. Microbiome 6:186.

Woodcock, D. J., P. Krusche, N. J. C. Strachan, K. J. Forbes, F. M. Cohan, G. Méric, and S. K. Sheppard. 2017. Genomic plasticity and rapid host switching can promote the evolution of generalism: a case study in the zoonotic pathogen Campylobacter. Sci Rep 7:9650.

Youngblut, N. D., G. H. Reischer, W. Walters, N. Schuster, C. Walzer, G. Stalder, R. E. Ley, and A. H. Farnleitner. 2019. Host diet and evolutionary history explain different aspects of gut microbiome diversity among vertebrate clades. Nat Commun 10:2200.

Zuo, J., L. Liu, P. Xiao, Z. Xu, D. M. Wilkinson, H. Grossart, H. Chen, and J. Yang. 2023. Patterns of bacterial generalists and specialists in lakes and reservoirs along a latitudinal gradient. Global Ecol Biogeogr 32:2017–2032.

