## Supplemental Figures for "Increased host diversity limits bacterial generalism but may promote microbe-microbe interactions"

**Supplemental Information**

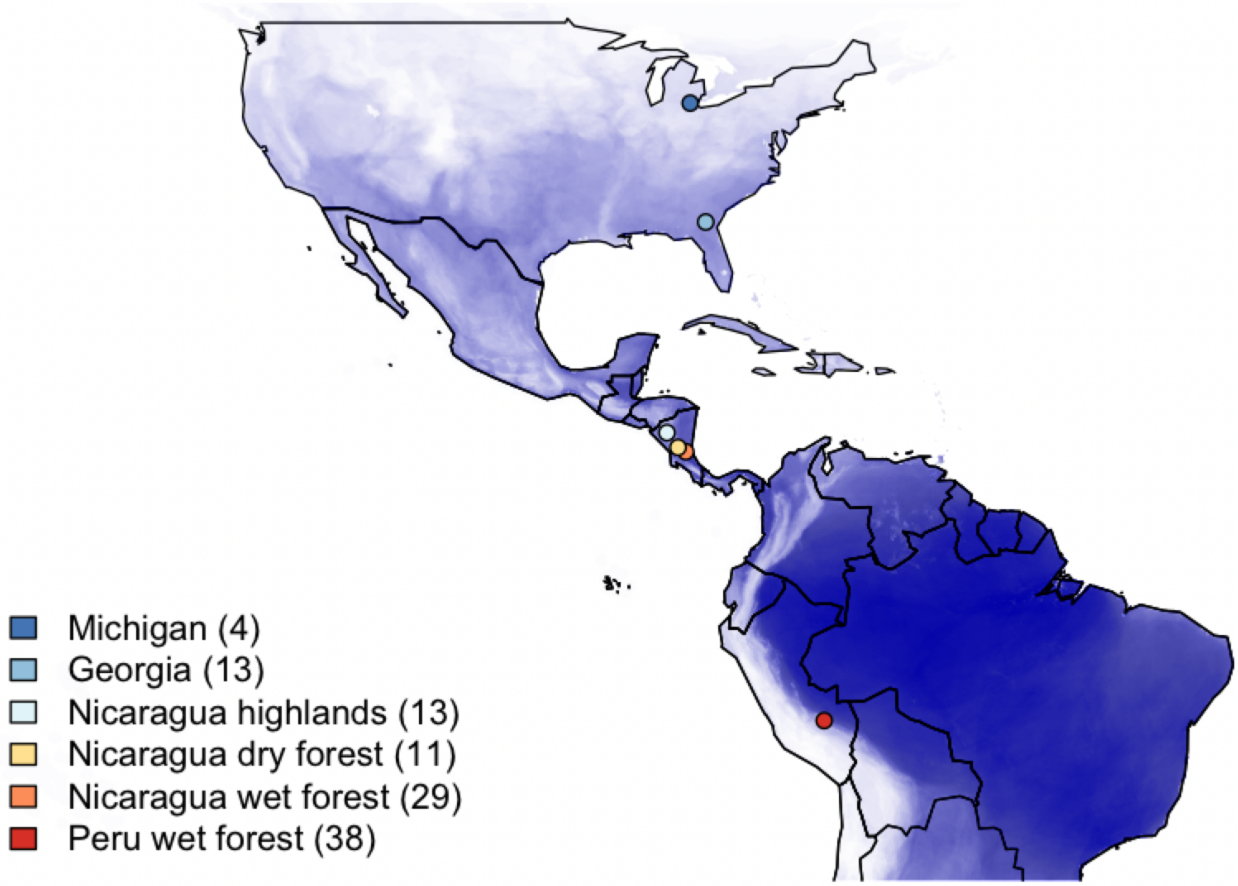


Figure S3: Map of collection locations overlaid on scaled reptile species richness from IUCN range polygons.

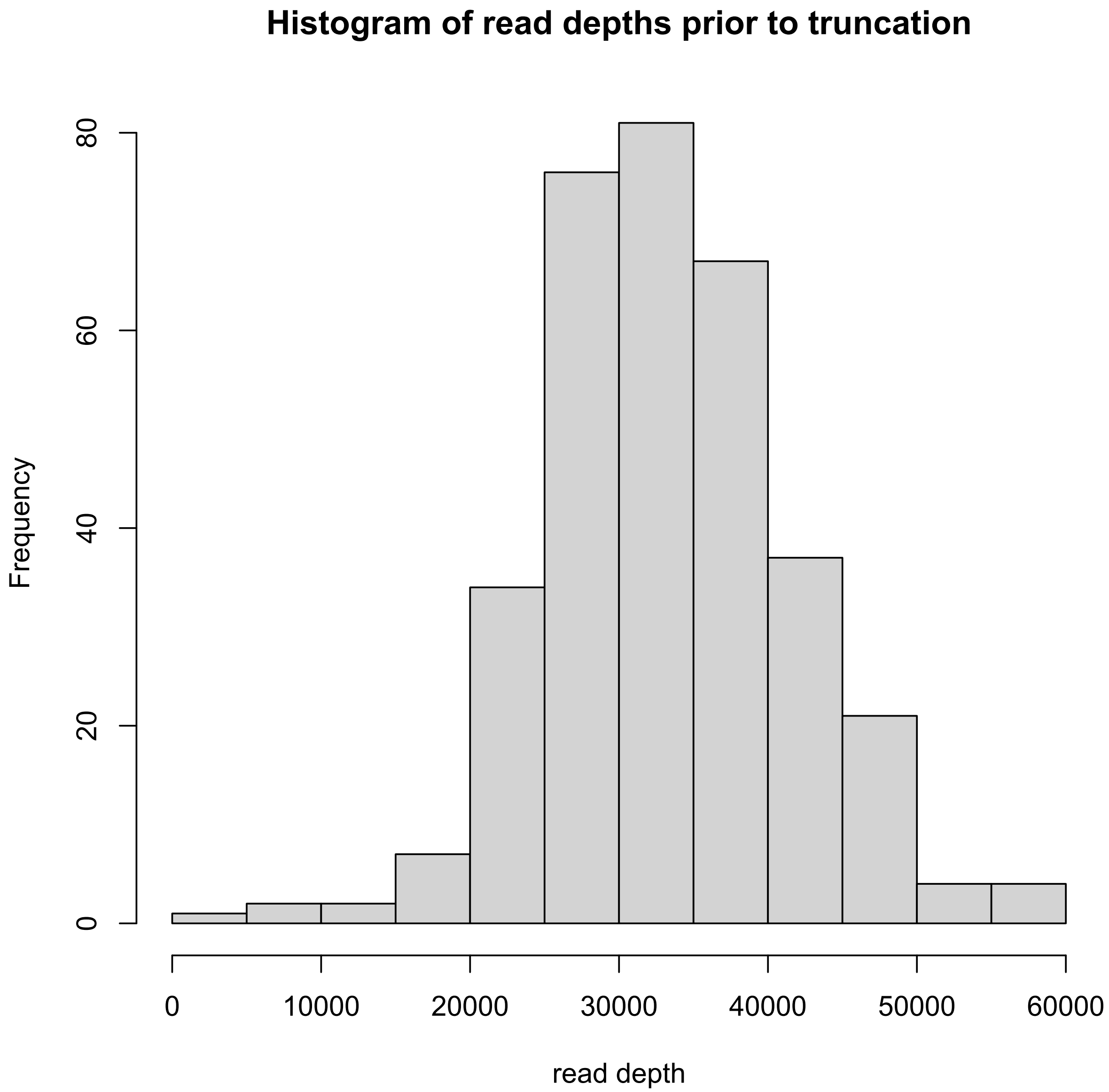


Figure S2: Read depths for all sampled hosts in our 16S rRNA dataset. For our analyses, we removed all samples with fewer than 10000 reads.

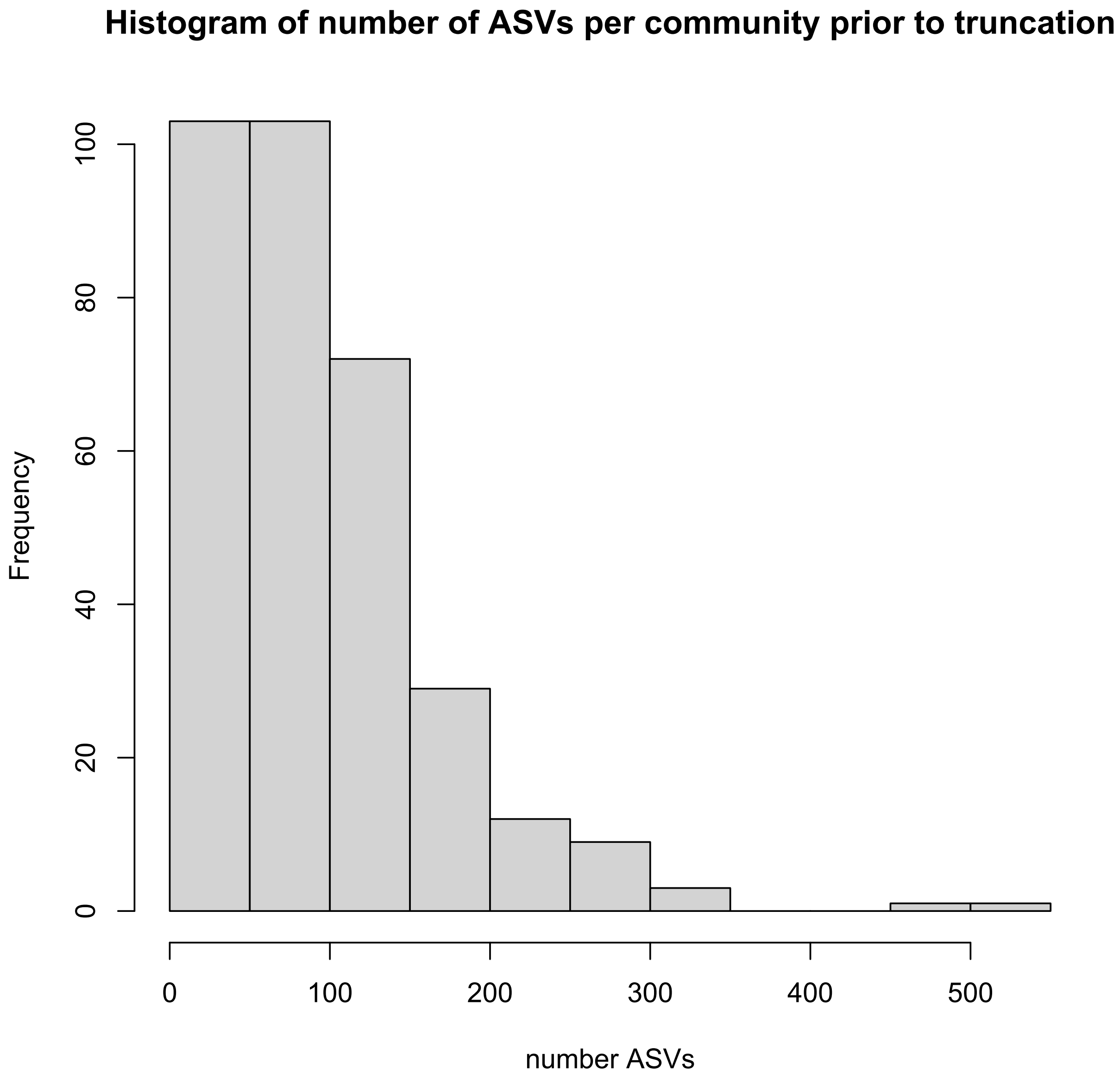


Figure S3: Number of ASVs per sampled host. We removed all sampled hosts with fewer than 50 ASVs.
